# SynPROTAC: synthesizable PROTACs design through synthesis constrained generative model and reinforcement learning

**DOI:** 10.64898/2025.12.10.693572

**Authors:** Mingyuan Xu, Qirui Deng, Hao Zhang, Anjie Qiao, Zhen Wang, Chang-Yu Hsieh, Hongming Chen, Jinping Lei

## Abstract

Protein hydrolysis targeting chimeric (PROTAC) has emerged as a promising technology in degrading disease-related proteins for drug design. Recent deep generative models can accelerate PROTAC design, but the generated molecules are often difficult to synthesize. Here we develop SynPROTAC model, which employs Graphormer encoded warhead or E3 ligand as input, and autoregressively samples reaction templates and building blocks through transformer based decoder for PROTAC construction. The model is also fine-tuned via reinforcement learning for generating PROTACs with favorable binding properties. The comprehensive evaluations indicated that SynPROTAC is capable of generating novel PROTACs with feasible synthetic routes, reasonable physico-chemical and binding related properties.

## Introduction

Protein hydrolysis targeting chimeric (PROTAC) has emerged as an effective tool for drug discovery since 2001. The rational design of PROTAC involves designing of three components including a warhead binding to POI, an E3 ligand binding to E3 ubiquitin ligase, and the linker between them^1-3^. Thus, compared to the design of conventional small molecular inhibitor, the design of PROTAC possesses multiple challenges^4^. Especially, design of PROTAC linker is particular challenging, because the linker segment is crucial for maintaining the conformational stability of ternary complex and associated with the chemical, pharmacokinetic properties and degradation efficiency of the PROTAC^5^.

Recently, deep learning based generative models have been developed and applied on PROTAC linker design^1,2,6-8^. For example, the previously developed SyntaLinker^9^ and PROTAC-RL^3^ utilize a transformer architecture to treat fragment linking as a sentence completion task using the 1D SMILES based representation. Recently, Li *et al*.^10^ developed DiffPROTAC which employs graph neural network (GNN) and transformer for generating new PROTAC linkers. Very recently, we developed a 3D generative model PROTAC-INVENT^5^, which generates both the PROTAC and its 3D putative binding conformation, and it is jointly trained with reinforcement learning (RL) to generate PROTAC with specified 2D and 3D based properties. However, all these generative models do not consider synthesizability of the generated PROTAC molecules in the generation process.

To overcome these challenges, we propose a generative model SynPROTAC that employs chemical reaction templates (RTs) and building blocks (BBs) to assemble molecules, and also incorporates RL to generate structurally novel, synthesizable PROTACs with desired binding properties. The synthetic route based generative model employs Graphormer encoded POI warhead or E3 ligand as input, and iteratively samples RTs and BBs through transformer based decoder under the guidance of RL for generating PROTACs with desirable binding properties. The performance evaluation of SynPROTAC revealed that it is able to produce novel PROTACs with high synthesis feasibility and desired 2D and 3D properties.

## Results

### Model Overview

As illustrated in Figure **1a**, the SynPROTAC model composes a chemical reaction path driven generative module for PROTAC generation along with synthetic route prediction, and a reinforcement learning module that navigates PROTAC generation towards desired binding related properties. The generative module is a transformer based network which employ Graph Transformer^11^encoded POI warhead or E3 ligand and reaction center embedding as input(Figure **1b**), and iteratively samples RTs and BBs for PROTAC construction (Figure **1c**). The synthetic pathway is a sequence consisting of three types of actions: selection of a BB or RT, and end of synthetic route. The action type in each step is predicted based on the sequence of actions generated so far and the reaction center enhanced molecular context of POI warhead and E3 ligand. The RTs and corresponding BBs are sequentially sampled based on the possibility distribution predicted by transformer in each step. Upon sampling the “end” action, the product PROTAC is generated by selecting a final RT to connect the remaining E3 ligand or POI warhead.

**Figure 1.**
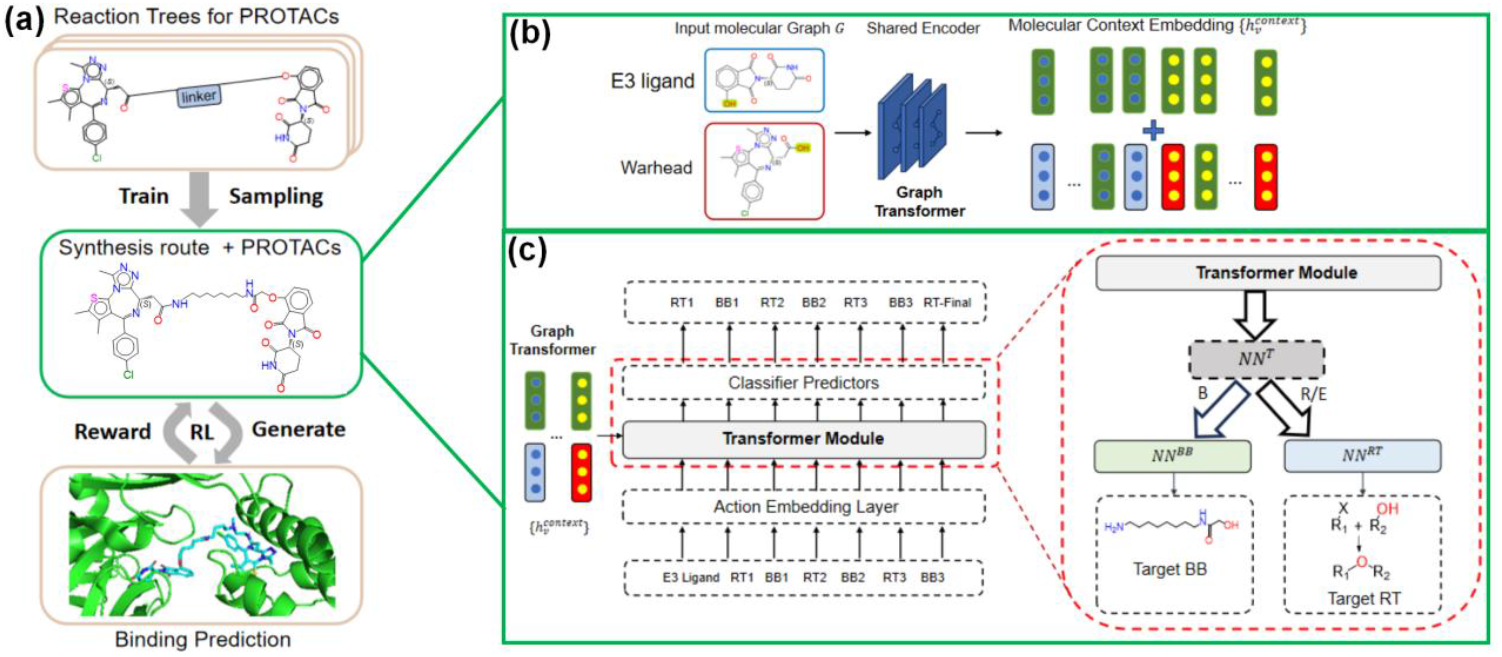
Overview of SynPROTAC model. (**a**) Overview of SynPROTAC model for design synthesizable PROTACs with desired properties. (**b**) The encoder block which employs a Graph Transformer to encode the input molecular graphs of both POI warhead and E3 ligand. The encoded atom-level embeddings are concatenated with embeddings of reaction sites as the molecular context controls. (**c**) The building blocks (BBs) and reaction templates (RTs) sampling block. The action embedding is firstly input into a transformed based decoder to predict the sampling action type token. If action type is B, the BB head predicts the building blocks; otherwise, *NN*^*RT*^ predicts reaction templates.

To train the synthetic path driven generative model for SynPROTAC, we firstly constructed a dataset of 20 million chemical reaction trees sampled from 91 RTs^12-14^ and 483 filtered BBs from Enamine database^15^ to meet the requirements for PROTAC constructions. The reaction trees were sampled by using MCTS^16-18^ algorithm based on the pairs of POI warhead and E3 ligand in PROTAC-DB 3.0 database^19^.

### Model Performance

The performance of SynPROTAC was evaluated and compared with the real PROTACs in PROTAC-DB 3.0 database^19^ on a range of metrics related with physico-chemical properties and synthesis feasibility^20^ score (Figure **2**). We selected a dataset of PROTACs with 608 pairs of ligands targeting POI and E3 ligase from PROTAC-DB 3.0 as the reference set, and 100 PROTACs were generated by SynPROTAC for each pair to perform assessment.

**Figure 2.**
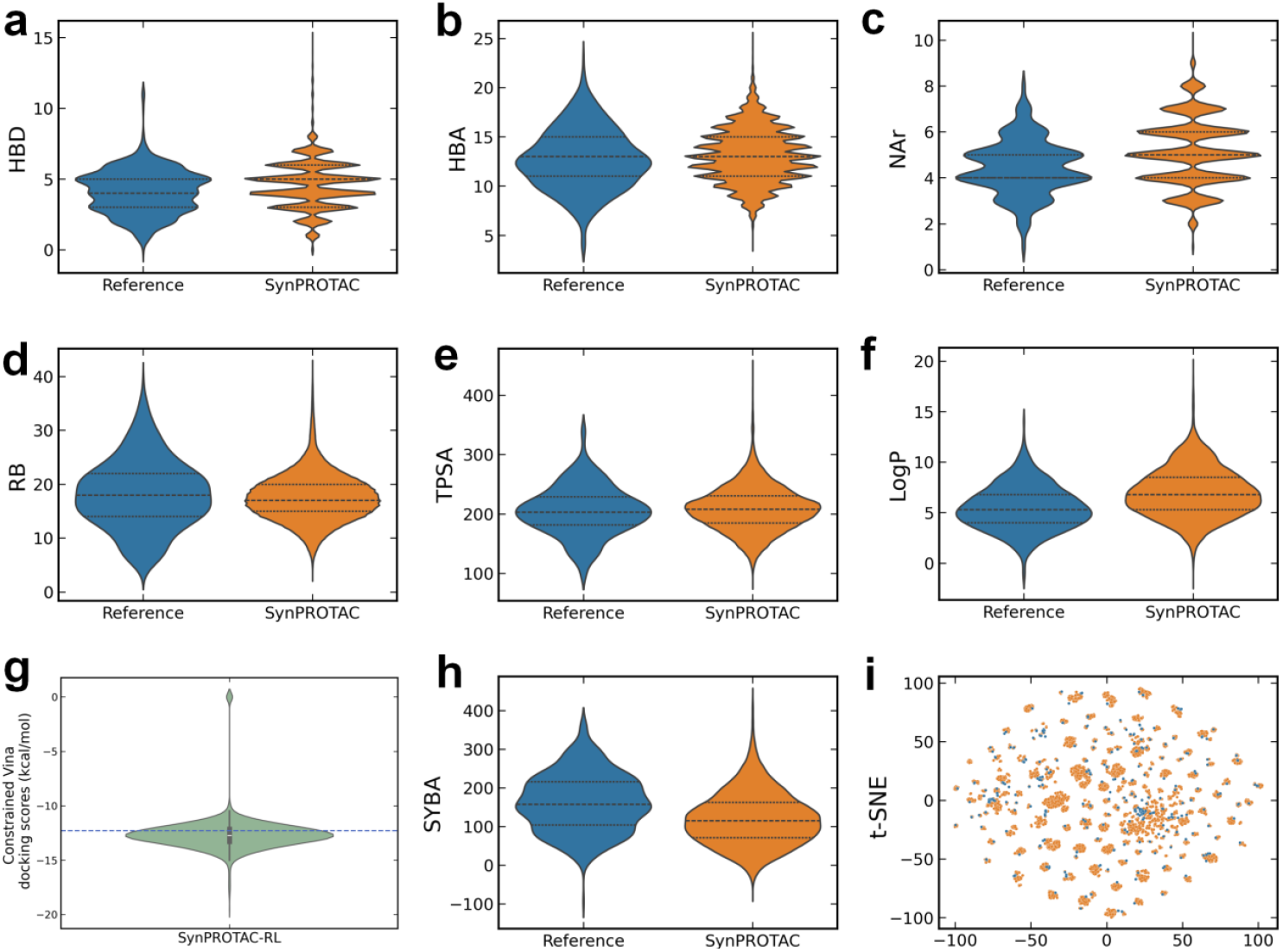
Performance of SynPROTAC. **(a-h**) The distributions of hydrogen bond donor (HBD), hydrogen bond acceptor (HBA), aromatic ring (NAr), rotatable bond (RB), topology polar surface area (TPSA), LogP (octanol-water partition coefficient), constrained AutoDock Vina docking scores, and synthetic bayesian accessibility (SYBA) score of PROTACs generated by SynPROTAC and the reference PROTAC set. (**i**) The t-distributed stochastic neighbour embedding (t-SNE) visualization of the reference and SynPROTAC set.

The physico-chemical properties of SynPROTAC set are as good as those reference ligands from PROTAC-DB (Figure **2a-f**). For example, the PROTACs assembled by SynPROTAC exhibit similar distribution on hydrogen bond donor (HBD), hydrogen bond acceptor (HBA), aromatic ring (NAr), rotatable bond (RB), and topology polar surface area (TPSA) to the reference molecules. Interestingly, PROTACs generated by SynPROTAC show larger average LogP (octanol-water partition coefficient) than reference set, indicating molecules generated by SynPROTAC exhibit larger lipophilicity and cell membrane permeability.

The AutoDock Vina docking score was used for estimating binding affinity of PROTAC with the POI/E3 complex. Here similar constrained docking protocol proposed in PROTAC-INVENT^5^ was adopted in the RL of SynPROTAC. Results in Figure **2g** shows that SynPROTAC could generate PROTACs with more favorable binding score than reference set, as the average docking score for SynPROTAC set is larger than reference set (-12.6 kcal/mol *vs* -12.3 kcal/mol).

The synthesizable properties (Figure **2h**) revealed that SynPROTAC enables generating PROTACs with high synthetic feasibility as reference molecules, and most of the molecules generated by SynPROTAC exhibit synthetic bayesian accessibility (SYBA) scores larger than 0, indicating PROTACs produced by SynPROTAC are easy to synthesize^20^. The t-SNE plot of chemical space distribution (Figure **2i**) revealed that SynPROTAC generated molecules covers significantly much larger space than reference set.

## Conclusion

Here, we developed SynPROTAC model, which employs transformer based generative alogrithm to iteratively sample chemical reaction templates and building blocks composing the synthesis routes of a PROTAC molecule. The model is further fine-tuned by RL to optimize the binding properties of generated PROTACs. These strategies enable SynPROTAC to generate novel PROTACs with high synthesis feasibility, favorable physico-chemical and binding-related properties, and make SynPROTAC potentially a useful tool for designing PROTACs in drug discovery projects.

## Methods

### Synthetic route based generative model

SynPROTAC adopts an encoder-decoder architecture, which includes a Graph Transformer-based^11,21^ encoder and a Transformer decoder. The synthetic route is represented as a sequence of actions in SynPROTAC. There are three types of actions to choose at each time step *t:* (1) [*B*], the selection of a BB, (2) [*R*], the selection of a RT; (3) [*E*], end of the synthesis route by selecting a RT to connect the remaining POI warhead or E3 ligand.

#### Graph encoder

The given pair of POI warhead and E3 ligand are represented as 2D molecular graph *G* = {ℋ, ℰ}, including the node features ℋ = {*h*_*v*_}for all nodes ∀*v* ∈ *V*, edge features ℰ = {*e*_*v,w*_}for all bonded pairs ∀(*v, w*) ∈ ℬ. The Graph Transformer denoted as *GE* is employed to encode the molecular graph. Each graph is encoded into a set of node-level representations 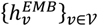 along with a padding mask through several graph attention layers with relative positional encodings as shown in Eq 1.

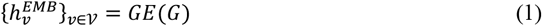

The embedding of reaction centers in both POI warhead and E3 ligand (predefined by users) are identified, and their representations are concatenated with all node features. As shown in Eq 2, the encodings of POI warhead and E3 ligand are merged via a MLP layer followed by layer normalization, forming a unified molecular context 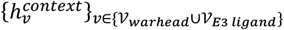.

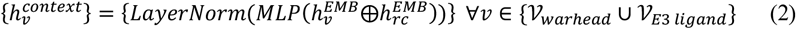

#### Synthetic action decoder

SynPROTAC employs a multi-layer Transformer to autoregressively predict synthetic actions. The Transformer takes the encoded molecular context 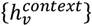 as input and generates a sequence of actions. The decoder utilizes masked self-attention to capture dependencies among previous actions and employs cross-attention to incorporate molecular context.

For a given synthetic route *A* = {*a*_*i*_}, action *a*_*i*_ is represented as the one-hot vector of reaction types *T*_*RT*_ if action type 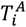 is [*R*] or [*E*]. Otherwise, *a*_*i*_ is the one-hot vector of building block index *T*^*BB*^. The synthetic action decoder introduces an action embedding layer, which contains several MLPs specified by different action type to encode {*a*_*i*_} into action embeddings 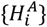 as shown in Eq 3.

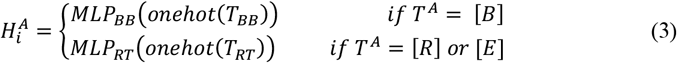

The Transformer module receives the action embeddings and generates embedding of next synthetic action autoregressively. To improve performance of Transformer module, sinusoidal positional encodings are added to the embeddings to indicate token order.

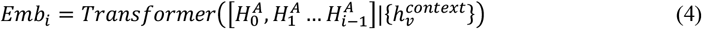

Three dedicated classifier predictors, including action type predictor *NN*^*T*^, building block predictor *NN*^*BB*^ and reaction template predictor *NN*^*RT*^ map the decoded embedding *Emb*_*i*_ to discrete outputs. For the *i*^*th*^ synthetic action step, *NN*^*T*^ predicts possibility distribution of action type 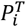 with *Emb*_*i*_. Then, the synthetic action type 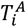 is predicted after *argmax* and *softmax* operations.^22^

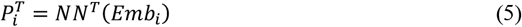

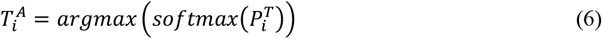

If 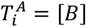, *NN*^*BB*^ predicts the possibility of suitable building blocks in 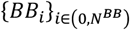, otherwise *NN*^*RT*^ predicts the reaction template *RT*_*i*_. To avoid the unapplicable RT or BB being sampled from previous action, a vector from the predefined BB and RT relation matrix *C*_*mask*_ is multiplied to the predicted possibility distribution:

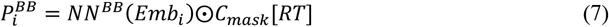

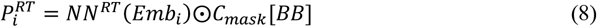

After *softmax* and *argmax* operation, the selected BB or RT is shown in Eq (9) ∼ (10):

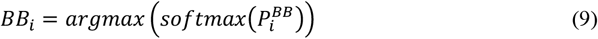

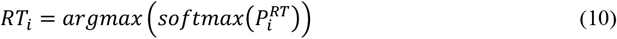

#### Model training

In the training phase, each synthetic action in the reaction tree is converted to action sequence {action type token *T*, action *a*} pairs. The training objective is shown in Eq 11, consisting of three components: 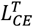 for action type prediction, 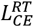 for reaction template prediction, and 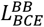 for Morgan fingerprint prediction of building blocks.

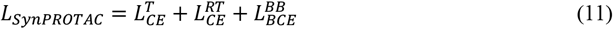

For a given synthetic route with length of *l*, the above loss terms can be illustrated in Eq 12∼14.

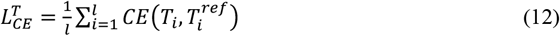

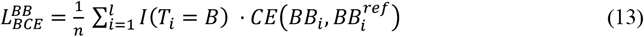

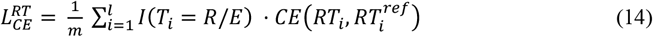

where *I* is the indicator function, *CE* represents cross entropy, *n* is the number of “*B*” tokens in the synthetic route, and *m* is the number of “*R*” and “*E*” tokens in the synthetic route.

### Reinforcement learning

The prior model generates molecules by iteratively sampling synthetic actions based on the input POI warhead and E3 ligand. To generate PROTAC molecules with desirable 2D/3D based properties, we integrated the prior model into a policy-based reinforcement learning workflow for molecular optimization. Here, we employ the similar RL algorithm proposed by Olivecrona *et al*.^*23*^ to maximize the reward of generated reaction path.

#### Binding conformation of PROTACs

For predicting 3D binding conformations of PROTACs and scoring them, we utilized a similar constrained docking workflow as our previously reported PROTAC-INVENT model^5^. This method takes the generated PROTAC linker, a pair of E3 ligand and POI warhead, and a reference PROTAC ternary structure (PTS) as input; then returns an initial 3D conformation of the complete PROTAC compound at the reference PTS binding site by doing a constrained force field minimization. Finally, the AutoDock Vina^24,25^ docking is carried out to further optimize docking poses of generated PROTACs and the docking scores are obtained.

#### Scoring of PROTACs

The AutoDock Vina docking score of the generated PROTAC molecules is used as the only reward component of RL:

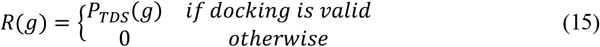

*P*_*TDS*_ refers to a transformed docking score which scales the original docking score to the range from 0 to 1.

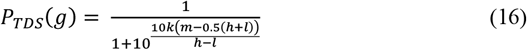

where *k* is a scalar, *m* represents the AutoDock Vina score value, *h* is the user-defined upper threshold of *m*, and *l* is the user-defined lower threshold of *m*.

## Notes

### Competing Interest Statement

The authors have declared no competing interest.

